# Revealing parasitoid-pest interactions using multi-marker metabarcoding: a nationwide assessment of parasitism in cabbage stem flea beetle larvae

**DOI:** 10.64898/2026.10.09.757974

**Authors:** Patricia A. Ortega-Ramos, Jordan P. Cuff, Darren M. Evans, Larissa Collins, Alice L. Mauchline, Robbie D. Girling, Samantha M. Cook

## Abstract

Management of insect pests of oilseed rape (*Brassica napus* L.) such as cabbage stem flea beetle (CSFB; *Psylliodes chrysocephala* L.) is increasingly challenging given a lack of effective permitted insecticides. Conservation biocontrol, including through naturally-occurring parasitoids of CSFB, is one of the first lines of defense and the baseline for integrated pest management (IPM) strategies, but we lack a comprehensive understanding of the parasitoids involved and how parasitism rates vary regionally. Such data have traditionally required rearing of parasitoids from their pest hosts, which is highly error-prone, laborious and overlooks instances of multiple parasitism and hyperparasitism.

Using multi-marker DNA metabarcoding, including a novel host-exclusion parasitoid PCR primer pair, we assessed parasitism of 1,040 CSFB larvae from 13 counties in England, UK. We confidently identified seven parasitoids of CSFB larvae, with ten taxa present in unfiltered data. Parasitism rates differed between regions, with the north of England having significantly lower parasitism rates than southern regions. Our novel beetle parasitoid PCR primers demonstrate a high congruence with visual detections of parasitoids and a high rate of novel interaction detections.

Ultimately, by determining both rates of parasitism and natural enemy identities, these data significantly extend our understanding of in-field CSFB parasitoid community dynamics. This information could guide the enhancement of farm management, especially using network science in decision support systems related to insecticide application, but also more broadly in the context of crop and habitat management for improved conservation biocontrol and IPM.

## Introduction

The cabbage stem flea beetle (CSFB, *Psylliodes chrysocephala* L.) is the most abundant and widely distributed pest of autumn-sown oilseed rape (OSR, *Brassica napus* L.) in Europe (Zheng et al., 2020). Both adult and larval stages cause severe damage to OSR: adult feeding threatens crop establishment during autumn, while larvae mine petioles and stems, reducing plant vigour, survival, and ultimately yield (Bonnemaison & Jourdheuil, 1954; Evans, 2007; Green, 2008; Williams & Carden, 1961). Historically, this pest has been controlled using synthetic insecticides, primarily neonicotinoid seed treatments and foliar pyrethroid applications; however, following the 2013 EU restriction on neonicotinoid seed treatments for OSR (European Commission, 2013), pyrethroid sprays have become the only remaining chemical control option (Ortega-Ramos et al., 2022). This heavy reliance on a single insecticide class has imposed intense selection pressure, resulting in widespread pyrethroid resistance across Europe (Heimbach & Müller, 2012; Højland et al., 2015; Willis et al., 2020; Zimmer et al., 2014). Consequently, control failures have led many farmers to abandon OSR cultivation, causing a sharp decline in OSR cultivation across the EU and UK (Andert & Ziesemer, 2021; Ortega-Ramos et al., 2022), with estimated annual economic losses of approximately £1 billion to the UK alone due to reduced production and increased imports (AHDB, 2025; United Oilseeds, 2024). To maintain sustainable production of this important crop, there is a crucial need for alternative and environmentally benign solutions as part of an integrated pest management (IPM) strategy.

Biological control by natural enemies represents a key ecosystem service that reduces crop losses from insect pests and constitutes an essential component of IPM (Naranjo et al., 2015). Biological control relies on ecological precision, understanding the parasitoids ecological needs and behaviors allow us to exploit their potential and better reduce the pest population. Therefore, resolving the identity and relative importance of these parasitoids is a prerequisite for conserving and exploiting them within IPM. Reliable and rapid detection would make it possible to measure the strength of this natural control and factor it into management decisions — informing conservation measures that support the parasitoid species contributing most to pest control, and guiding spray thresholds that avoid insecticide use where parasitism is already suppressing CSFB populations.

The CSFB is attacked by various natural enemies, including seven known species of parasitic wasps reported to parasitise CSFB larvae (Ulber et al., 2010). Among these, *Tersilochus microgaster* Szépligeti, 1899 has shown strong potential as a biological control agent against CSFB, with field parasitism rates of 24.6%, 44.4%, and 22.8% recorded in Germany during 2001– 2003, respectively, at peak host larval abundance in early May (Ulber & Nitzsche, 2006). In the UK, Ferguson et al., (2006) documented *T. microgaster* with an 11 % larval parasitism rate in 1999 and Barari, et al., (2005) recorded 7.7 % parasitism in 2002. *Tersilochus tripartitus* Brischke, 1880 was long considered the most common and widespread parasitoid of larval stages of CSFB in spring, with parasitism rates ranging from 3 to 60% (Aubert & Jourdheuil, 1958; Dosse, 1961; Ulber, B. & Williams, 2003); however, Ulber et al. (2010) suggested that many historical *T. tripartitus* records were likely misidentifications of *T. microgaster*, obscuring the relative importance of both species. Other documented parasitoids include *Cremastus carinifer* Thomson, *Aneuclis melanarius* Holmgren, *Diospilus oleraceus* Haliday, *Diospilus morosus* Reinhardt and *Trichomalus lucidus* Walker, though these are considered of minor importance, with parasitism rates below 6% or lacking quantitative data (Ulber & Wedemeyer, 2004; Ulber et al., 2010).

Parasitism in OSR and a range of other systems has traditionally been assessed through rearing and/or dissection of hosts, followed by taxonomical identification; rearing is technically demanding, time-consuming, space-intensive, and error-prone (Barari et al., 2006). In addition, identification of parasitoids based on morphological characters is challenging and, although some keys are available for parasitoids of oilseed rape pests (Alford, 2003; Ferguson et al., 2010), misidentifications are common due to their small size, ambiguous key characteristics and the presence of cryptic species and phenotypic variation within populations (Atanassova et al., 1998). Molecular approaches offer potential to overcome these limitations Miller et al., 2021; Sow et al., 2019), enabling standardized, rapid, and cost-effective identification of ecological interactions including host-parasitoid associations (Hrček & Godfray, 2015; Šigut et al., 2017).

Molecular diagnostic approaches, such as diagnostic PCR and loop-mediated isothermal amplification, can rapidly verify the presence of given species within a sample (Gariepy et al., 2007; Miller et al., 2025, 2026; Staudacher et al., 2016); however, this approach is limited to known interactions with specific target taxa, ignoring the potential for previously undocumented associations (Rennstam Rubbmark et al., 2019). DNA metabarcoding enables the simultaneous detection of multiple species from the same sample via high-throughput sequencing (Taberlet et al., 2012), enabling identification of diverse assemblages of morphologically cryptic and unexpected taxa (Gariepy et al., 2007). While this allows parasitoid identification even when host DNA is co-amplified, extensive host amplification can obscure or completely mask parasitoid signals (Cuff et al., 2023). Both diagnostic PCR and metabarcoding have successfully detected host-parasitoid interactions in previous studies (Farrokhzadeh et al., 2017; Kitson et al., 2018; Šigut et al., 2017), demonstrating their potential for resolving these complex ecological networks. Nevertheless, a high-throughput, taxonomically comprehensive, and efficient approach specifically designed for detecting parasitoids in beetle hosts remains lacking.

Here, we use multi-marker DNA metabarcoding to investigate host-parasitoid interactions in CSFB populations across the UK. To achieve this, we (1) designed novel PCR primers suitable for both diagnostic PCR detection and DNA metabarcoding of endoparasitoids in CSFB; (2) compared rated of parasitism using DNA metabarcoding between general and exclusion PCR primers, and with traditional visual assessment approaches; and (3) assessed parasitoid community composition and prevalence across UK regional populations of CSFB to determine differences in biocontrol pressure and natural enemy diversity. The approaches developed for this study show promise for broader application to host-parasitoid interactions, and the data generated provide insight into the community dynamics of CSFB, with potential to inform agricultural management practices for these prolific crop pests.

## Methods

### Sample collection

Plants collected in spring (mid-March) 2019 as part of a Defra-funded OSR pest survey from 33 different fields across seven regions in England (see below for details) were dissected manually to extract any CSFB larvae present. Twenty-five plants were randomly sampled from each field. Petioles and main stems were dissected under sterile conditions with a binocular microscope, using a scalpel and the number of CSFB larvae recorded. Larvae were individually kept in microcentrifuge tubes containing 95% ethanol and labelled according to the site, plant number and the petiole number where the larvae were found. The scalpel blade was disinfected (rinsed with bleach) after each larva was found, and blades were changed between plants. To ensure that even low prevalence parasitism was detected (expected to be between 0 and 10%), only sites with high larval numbers (>125 larvae per 25 plants) were considered in the study (n = 13, North East (1), Yorkshire (2), East Midlands (1), West Midlands (1), East (3), South East (3), South West (2). To facilitate the analysis we used the Fera region assignment (Collins 2017) (North = North East + Yorkshire; East = East + East Midlands, Midlands & West = West Midlands, South East = South East; South West = South West). In total, 3757 larvae were extracted. On average, 268.4 larvae were found per field across 25 plants.

### Visual assessment of parasitism

Before beginning DNA extraction, larvae were visually assessed for parasitism by manual dissection under a binocular microscope. Larvae were considered parasitised if parasitoid eggs or larvae were visibly present within the host body cavity (Figure 1).

**Figure 1:**
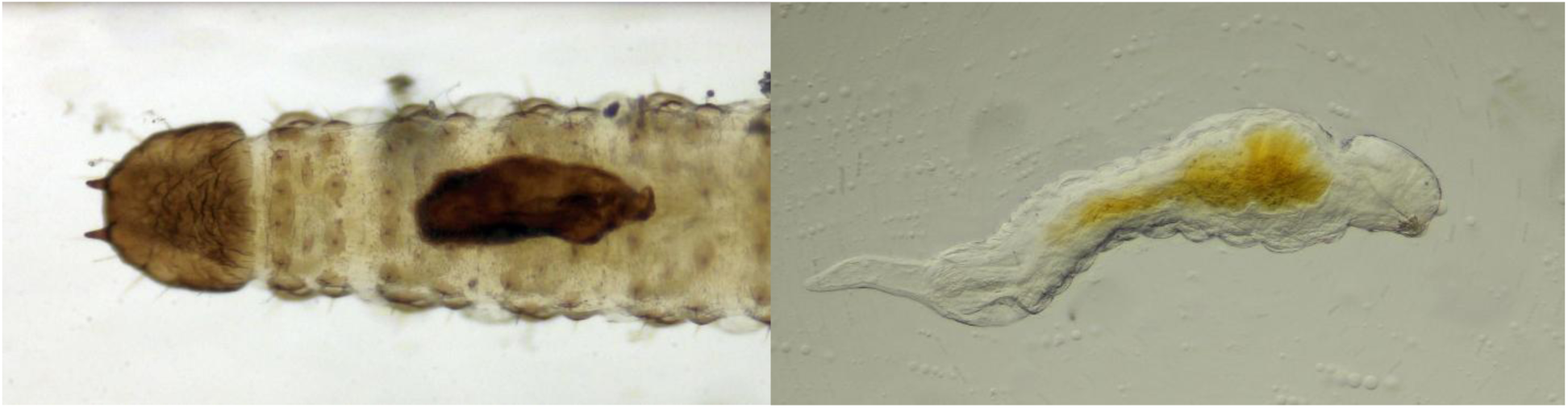
Examples of a visually assessed parasitoid egg (left) and larva (right) from within cabbage stem flea beetle larvae.

### DNA extraction

DNA was extracted in a 96-well plate format using a modified BOMB-Bio total nucleic acid extraction protocol (Oberacker et al., 2019). The CSFB larvae were dissected ventrally to facilitate the digestion of the inside of the larvae and to extract more effectively the parasitoid’s DNA. Dissections were made with a scalpel and forceps that were cleaned with Chemgene® and distilled water between every individual dissection. Each larva was placed in a 1.2 ml tube and left in a heat block for 30 min to ensure all ethanol was fully evaporated. Buffer (60 µl TNES buffer comprised of 20 mM EDTA, 400 mM NaCl and 50 mM Tris) was added to each larva, along with 2 µl Proteinase K and 3 µl RNAse A. The larva was then incubated in this mixture overnight at 55 °C. The resulting supernatant was mixed with 120 µl 1.5X GITC lysis buffer (6 M GITC, 75 mM Tris-HCl, 3% sarkosyl, 30 mM EDTA, 0.15% antifoam) and 240 µl isopropanol, and shaken at room temperature at 1400 rpm for 5 min. Carboxyl-coated magnetic beads (120 µl TE-diluted) were added to the cell lysate. The beads were settled on a magnetic stand; the supernatant was discarded and 400 µl isopropanol added and mixed. Magnetic beads were settled again to discard the supernatant and washed twice with 300 µl of 80% ethanol. After washing, the supernatant was removed completely, and the beads were dried on a heating block until completely dry (∼30 min). The extracted DNA was resuspended in 70 µl of nuclease-free water for PCR amplification.

### PCR primer design

Existing metabarcoding PCR primers were tested against CSFB and parasitoid samples, but amplification of DNA extracted solely from CSFB indicated it was likely that large proportions of data output would be lost to CSFB, and the results would not be diagnostic without subsequent high-throughput sequencing. Novel PCR primers were therefore designed to match hymenopteran parasitoids without amplifying beetle DNA (henceforth, beetle exclusion parasitoid primers). Mass alignments of COI sequences were batch-downloaded from (Ratnasingham & Hebert, 2007) GenBank (Benson et al., 2013) and BOLD (Ratnasingham et al., 2024) using PrimerMiner (Elbrecht & Leese, 2016) in R v.3.3.4 (R Core Team, 2020) to facilitate visual inspection of existing and novel primer sites. PrimerMiner clusters batch downloads into operational taxonomic units (OTUs) based on sequence similarity and visualises mass alignments of sequence data for primer design. By merging overrepresented and duplicate sequences through taxonomy-independent clustering, PrimerMiner accounts for within-species variation and cryptic species whilst ignoring rare haplotypes (Elbrecht & Leese, 2016). Sequences were downloaded for all terrestrial invertebrate orders available in 2017 (when this alignment library was constructed), and consensus sequences were created by clustering these into OTUs for each order. The COI sequences were trimmed to include only the Folmer region (Folmer et al., 1994) using Geneious R10 (Kearse et al., 2012) for subsequent use in PrimerMiner. Alignments created via PrimerMiner included Hymenoptera, Coleoptera and Hemiptera (the latter included to ascertain any potential for these primers in the assessment of hemipteran parasitoids, e.g., aphid parasitoids), to find primer sites conserved across Hymenoptera, but different for Coleoptera. Sequences for CSFB specifically were also downloaded, aligned, and trimmed for direct comparison. Four viable primer sites were identified, and primers designed with degeneracies to facilitate broad amplification of Hymenoptera. Four novel primers were designed, different permutations of which amplify varying fragment sizes of the COI gene (Table 1, Figure 1). Primer melting temperature and GC content were determined using the online ThermoFisher Scientific Multiple Primer Analyzer tool.

**Table 1:**
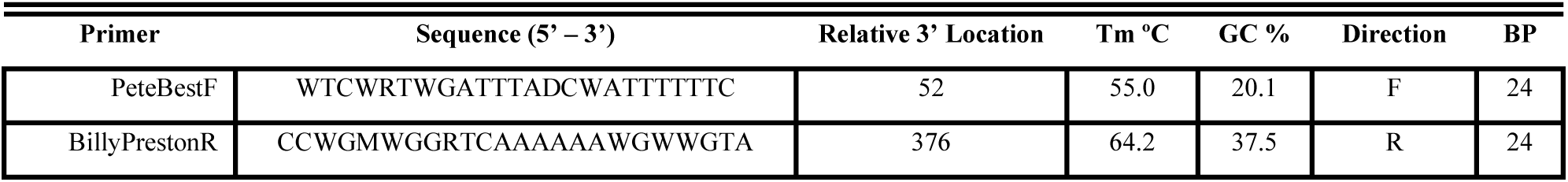
PCR primers designed within the COI gene for parasitoid amplification and beetle exclusion. Tm = melting temperature; GC % = guanine/cytosine content; BP = length in base pairs.

The coverage of primers (% amplified) was determined via PrimerMiner using the same mass alignments used for primer design. PrimerMiner uses a taxonomy-independent database and accounts for adjacent base mismatches and the position of each base in the primer. Viable pairings of novel primers, and one pairing including the general primer BerenF (Cuff et al., 2021), were compared against the mass alignments of Hymenoptera, Coleoptera and Hemiptera, and the available sequence data for CSFB. Given the lowest amplification of Coleoptera and highest amplification of Hymenoptera, the combination PeteBestF-BillyPrestonR was selected for further use.

The optimal primer pair was tested in vitro against DNA extracted from beetle and parasitoid tissue, and larval CSFB samples including two with visually confirmed parasitism. Primer testing took place in 15 µl reaction volumes comprised of 7.5 µl MyTaq Red Mix 2X (Bioline), 0.2 µM of each primer and 3 µl template DNA. Temperature gradient PCRs were used to determine the optimal annealing temperature for the selected primer pair, with temperatures between 50 °C and 60 °C considered. Optimal PCR conditions were ultimately determined to be: 95 °C for 1 min, 35 cycles of 95 °C, 51 °C and 72 °C for 30, 30 and 60 s, respectively, and a final extension at 72 °C for 5 minutes. Successful amplification was confirmed by gel electrophoresis.

### DNA metabarcoding

DNA was amplified using two PCR primer pairs: the general invertebrate primers miCOIintF and jgHCO2198 (Leray et al., 2013), and the newly designed beetle exclusion parasitoid primers PeteBestF and BillyPrestonR. This multi-marker approach was adopted to ensure a sufficient depth of detections (via exclusion primers) and a sufficient breadth of detections (via general primers; Cuff et al., 2023). PCR plates included a negative control (comprising molecular grade water instead of DNA) for each column and row, two positive controls (oak processionary moth extract) and two blanks (for which no reagents were added), leaving 80 samples per plate. PCR primers were tagged with unique 8 bp tags (Taberlet et al., 2018) to identify samples by their combination of forward and reverse tags following sequencing, and primer sequences contained Nextera bridge primer sites for PCR-based adapter attachment. PCR reactions were sealed with mineral oil to limit evaporation and cross-contamination (Kitson et al., 2019).

The 20 µL reactions comprised 10 µL MyTaq Red Mix, both forward and reverse primers at 0.2 µM, 4 µL DNA and the remaining volume made up with molecular grade water. Reactions for both primer pairs were carried out under the following temperature regime: initial denaturation at 95 °C for 1 min, followed by 35 cycles of 95 °C for 30 s, 51 °C for 30 s and 72 °C for 60 s, followed by a final extension at 72 °C for 5 min. Optimal annealing temperatures were determined by temperature gradient PCR. Amplification was confirmed by gel electrophoresis using 2% agarose gels.

PCR products were normalised to reduce variation in concentration across samples. This was performed using 0.1X carboxylated solid phase reversible immobilisation (SPRI) beads (Sera-Mag Magnetic SpeedBeads) at a 1:1 ratio of sample to bead solution. Samples were mixed with beads and incubated at room temperature for 5 min before being placed on a magnet for 5 min. All but 5 µL of the supernatant was removed and 200 µL 80% ethanol added to each sample. The ethanol was removed and replaced with a second volume of 200 µL 80%. This ethanol was removed, the samples were removed from the magnet, and the beads were allowed to dry for ∼ 30 s. To each well, 10 µL molecular grade water was added, and the solution mixed and incubated for 5 min. The beads were placed back on a magnet for 5 min, after which 8 µL of the supernatant from each sample was removed and pooled for each 96-well plate. Of this pooled sample, 700 µL was cleaned once more as above, but this time with 1X SPRI beads, eluting into 100 µL of molecular grade water.

The concentration of each pool was determined by Qubit high-sensitivity double-stranded DNA assay and diluted to approximately 5 ng µL^-1^. Preparation of libraries for sequencing followed the Nextera PCR route (Bohmann et al., 2022). A second round of PCR was carried out with Nextera PCR primers for the attachment of Illumina indexes, each plate receiving a unique combination of tags (this nested-tagging approach allows differentiation of plates and samples, respectively (Kitson et al., 2019). Reaction volumes of 15 µL comprised 7.5 µL MyTaq Red Mix, 1.5 µL of Nextera sequencing adapter primer mix, 5 µL DNA library and the remaining volume made up with molecular grade water. Reactions were, as above, sealed with mineral oil. PCR reactions were carried out under the following temperature regime: initial denaturation at 95 °C for 5 min, followed by 12 cycles of 95 °C for 30 s, 60 °C for 30 s and 72 °C for 1 min, followed by a final extension at 72 °C for 10 min. Library DNA concentrations were measured using Qubit high-sensitivity double-stranded DNA assay, libraries were purified, as above, using SPRI beads, but with a 0.8X bead to sample ratio to remove small non-target fragments, and correct ligation of adapters was confirmed via D1000 tape on a TapeStation 4200. Libraries were sequenced on an Illumina MiSeq (2 x 300) at Newcastle University’s Genomics Core Facility.

### Bioinformatics

Adapter and primer sequences were removed, and reads demultiplexed and assigned to samples, using cutadapt (Martin, 2011) with no insertions or deletions permitted in tag sequences and a maximum error rate of 15%. Sequences were filtered, trimmed, denoised and assigned to amplicon sequence variants (ASVs) using DADA2 (Callahan et al., 2016). Taxonomic identities were assigned to ASVs via BLASTn with a custom database of animal COI sequence data extracted from both GenBank (Benson et al., 2013) and BOLD (Ratnasingham et al., 2024). The assignment with the highest bitscore was retained, and the taxonomic resolution of assignments was adjusted based on percent identity, with taxa being assigned to species, genus, family, order or class based on percentage identities of >97%, >95%, >90%, >80% and <80%, respectively. Samples with fewer than 1,000 reads following taxon assignment were removed. Sequences attributed to CSFB were removed given that these were detections of the host itself, a common problem in dietary metabarcoding (Cuff et al., 2023). Minimum sequence copy thresholds were used to remove potential false positives, with detections removed when the read count within a sample was less than the read count for that taxon in a negative control or blank; reads were otherwise removed using per-plate sample read count thresholds based on the maximum sample read count percentage comprised of suspected false positives (e.g., positive control taxa; (Drake et al., 2022)). All remaining read counts were converted to binary presence/absence data due to the error-prone semi-quantitative nature of read count data (Lamb et al., 2019). Non-parasitoid taxa were removed from the dataset and presumed to be surface contaminants, accidental consumption of plant tissue bearing that DNA or another route of introduction of non-target DNA.

### Data analysis

All analyses were performed in R 4.5.2 (R Core Team, 2025). Data were processed and transformed using the tidyverse package for transparency and reproducibility (Wickham et al., 2019). General and exclusion PCR primers were compared by determining their congruence with visual assessment data, their congruence with just samples deemed to be parasitised via visual assessment and just molecular detections that were not detected visually. To examine regional variation in CSFB parasitoid communities and parasitism rates across England, a generalized linear model (GLM) and a multivariate generalized linear model (MGLM) were run. The GLM compared parasitism status (parasitised/non-parasitised) for each individual CSFB larva between regions, with a binomial error distribution and logit link function. Model assumptions were checked using the ‘DHARMa’ package (Hartig, 2024) and post-hoc pairwise comparisons were made using the ‘emmeans’ package (Lenth & Piaskowski, 2025). Detected parasitoid communities were compared between regions using a multivariate generalized linear model (MGLM) using the ‘mvabund’ package (Wang et al., 2012), with a binomial error family, Monte Carlo resampling and adjusted univariate p values.

## Results

### Development and validation of parasitoid-specific PCR primers

Four novel PCR primers were designed within the mitochondrial COI gene to selectively amplify hymenopteran parasitoids while excluding coleopteran host DNA (Table 1). *In silico* evaluation of three primer pairings of these novel primers (PeteBestF-BillyPrestonR, PeteBestF-GeorgeMaRtin, and BrianFstein-BillyPrestonR), together with one pairing incorporating the general invertebrate primer BerenF (Cuff et al., 2021), revealed varying taxonomic selectivity (Figure 2; Figure S1). The primer pair PeteBestF-BillyPrestonR showed the highest predicted amplification of Hymenoptera (>90% of reference sequences) whilst also excluding CSFB and Coleoptera more generally (<5% of reference sequences; Figure 2).

**Figure 2:**
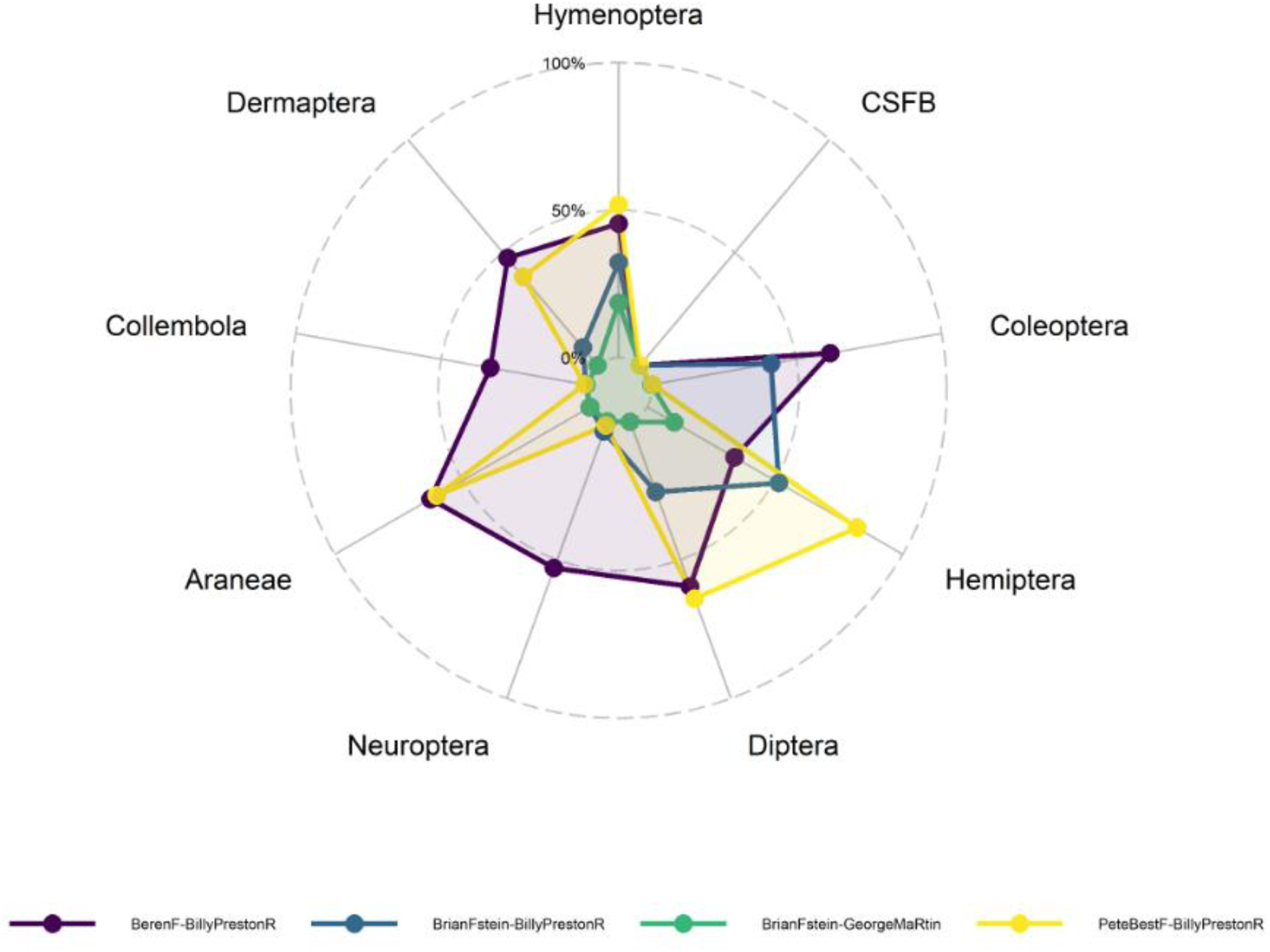
Exclusion PCR primers for the analysis of hymenopteran parasitoids in cabbage stem flea beetle (CSFB). The distance of each point from the center of the plot relates to the percentage of that group theoretically amplified by each primer pair, denoted by different colours.

*In vitro* validation confirmed the specificity predicted by *in silico* analysis PCR using PeteBestF-BillyPrestonR successfully amplified DNA from visually confirmed parasitised CSFB larvae, producing clear bands of the expected size (∼325 bp), while consistently failing to amplify DNA from non-parasitised beetle larvae (Figure S2). These results demonstrate that the primer pair effectively excludes beetle DNA while selectively amplifying parasitoid DNA present in mixed host-parasitoid samples.

### Molecular detection of parasitoids and comparison with visual assessment

Of the 1,040 CSFB larvae screened for parasitism across the 13 sites, visual assessment identified 62 (5.96%) larvae as parasitised, while metabarcoding identified 180 (17.31%) larvae as being parasitised (exclusion primers = 171, 16.44 %; general primers = 9, 0.87%) following minimum sequence copy threshold filtering. Prior to filtering, metabarcoding identified 633 (60.87%) larvae as being parasitised (exclusion primers = 598, 57.50%; general primers = 35, 3.37%).

The congruence between visual and molecular approaches for parasitoid detection varied markedly depending on the PCR primers used (Figure 3). When using exclusion primers, overall congruence between methods was low, at 24.8% (18.9% prior to filtering), while the use of general primers resulted in higher congruence of 92.8% (90.5% prior to filtering). This pattern was, however, reversed when examining detection accuracy for samples confirmed as visually positive for parasitism. For visually positive samples, exclusion primers demonstrated perfect congruence (100 %; 98 % unfiltered) whereas general primers showed poor agreement with visual detection for these known positive samples (3.57%; 3.77% unfiltered). Exclusion primers provided 88 detections not represented by the visual assessment (390 prior to filtering), whereas general primers provided 8 (33 prior to filtering).

**Figure 3:**
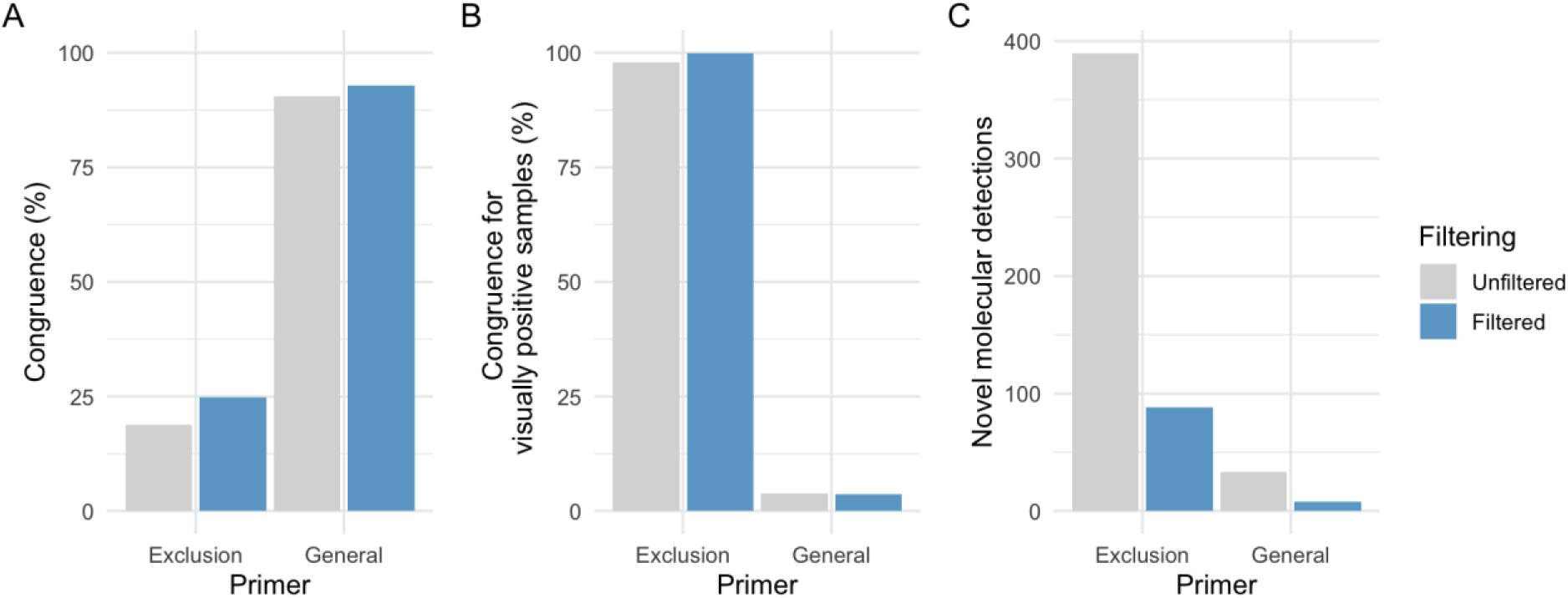
Congruence between visual and molecular approaches to parasitoid detection, with PCR primer pair and data filtering represented. A: Overall congruence of results, including both presence and absence of parasitoids. B: Congruence for just samples in which parasitoids were visually detected. C: Molecular detections in samples for which no parasitoids were visually detected.

### Parasitism rate

The overall parasitism rate across all studied regions of England was 11.5 %, varying substantially across sites (range: 1.25-23.75 %) and regions (range: 4.17-18.10 %). Region-level means were: East 12.5%, Midlands & West 16%, North 4.5%, South East 12% and South West 18%. The rate of parasitism, irrespective of parasitoid identity, differed significantly between regions of England (Table 2; Figure 4). Specifically, parasitism rates were significantly lower in the North compared to the East (GLM: estimate = 1.190 ± 0.365, z = 3.263, p = 0.010), the Midlands and West (GLM: estimate = 1.496 ± 0.443, z = 3.377, p = 0.007), the South East (GLM: estimate = 1.111 ± 0.38, z = 2.921, p = 0.029) and the South West (GLM: estimate = –1.628 ± 0.383, z = –4.253, p < 0.001). No other pairwise comparisons were significantly different (Table 2).

**Figure 4:**
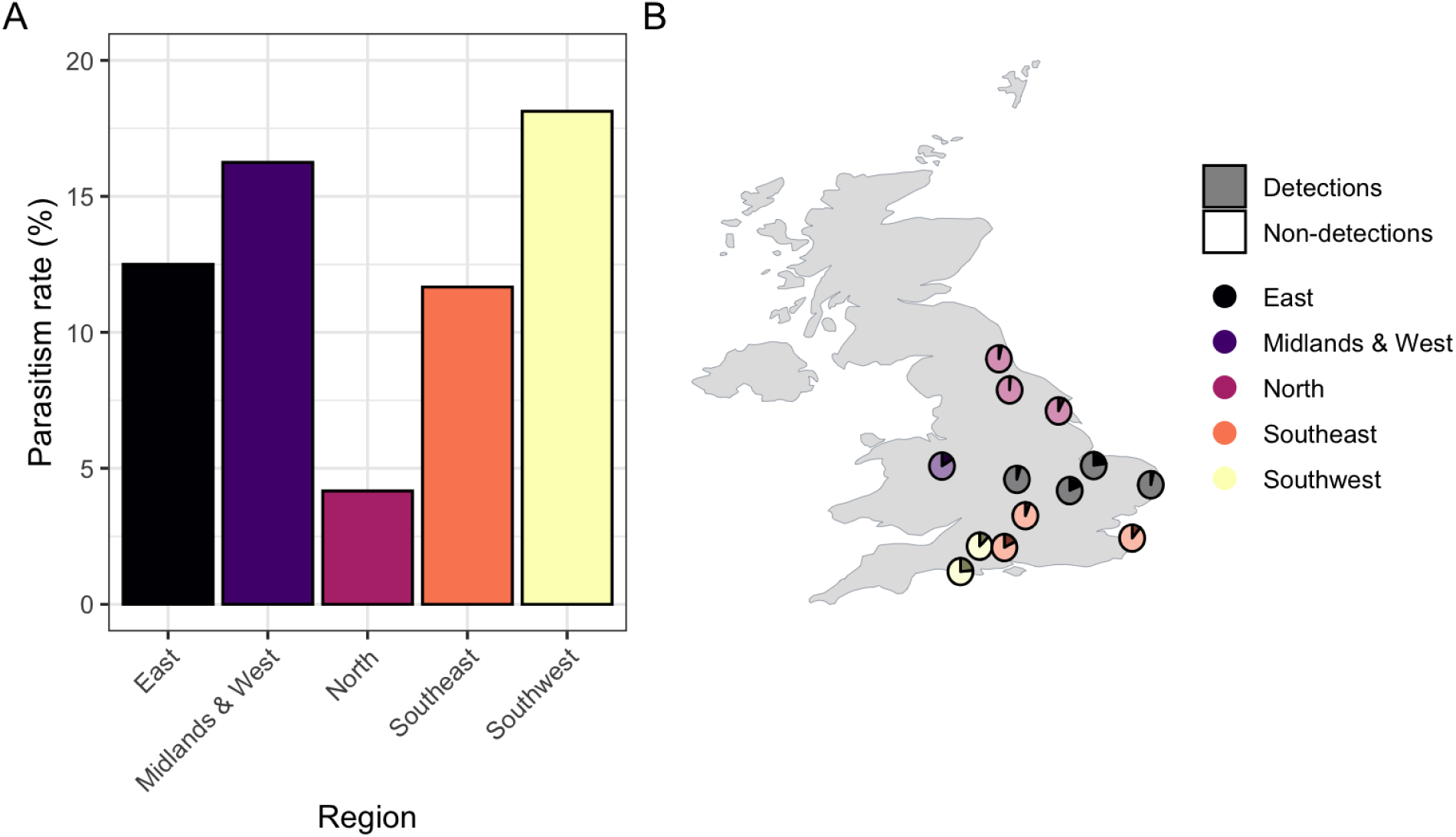
Parasitism occurrence rate of cabbage steam flea beetles (*Psylliodes chrysocephala*) across English regions and sites included in this study as a percentage of the beetles tested. A: Direct comparison of parasitism rates. B: Representation of parasitism rates at each site, coloured according to its corresponding region.

**Table 2:** Pairwise comparison of parasitism occurrence of cabbage stem flea beetles (*Psylliodes chrysocephala*) across regions of England from a binomial generalized linear model.

|  |  |  |  |  |
| --- | --- | --- | --- | --- |
|  | Midlands & West | North | South East | South West |
| East | estimate = $-0.306 \pm 0.347$, z = -0.882, p = 0.904 | estimate = $1.190 \pm 0.365$, z = 3.263, p = 0.010 | estimate = $0.078 \pm 0.263$, z = 0.299, p = 0.998 | estimate = $-0.438 \pm 0.266$, z = -1.647, p = 0.467 |
| Midlands & West | | estimate = $1.496 \pm 0.443$, z = 3.377, p = 0.007 | estimate = $0.385 \pm 0.364$, z = 1.058, p = 0.828 | estimate = $-0.132 \pm 0.366$, z = -0.360, p = 0.996 |
| North | | | estimate = $-1.111 \pm 0.38$, z = -2.921, p = 0.029 | estimate = $-1.628 \pm 0.383$, z = -4.253, p < 0.001 |
| South East | | | | estimate = $-0.516 \pm 0.287$, z = -1.798, p = 0.375 |

### Parasitoid community composition and occurrence across regions

After applying minimum sequence-copy thresholds and removing non-parasitoid detections, seven taxa were retained in the dataset (Figure 5). The most frequently detected was *Tersilochus microgaster* (Hymenoptera: Ichneumonidae; 164 detections, 64.3% of parasitised larvae), the detection frequency of which was nearly threefold that of the second most prevalent parasitoid (*Tersilochus* sp.; 59, 23.1%). *Tersilochus obscurator* was also highly prevalent (21, 8.2%). The remaining taxa were detected 1-6 times in total, and comprised *Tersilochus fulvipes*, Alysia sp. (Hymenoptera: Braconidae: Alysiinae), Pteromalidae sp. (Hymenoptera: Apocrita: Chalcidoidea) and the entomopathogenic nematode Rhabditida sp. (Nematoda: Chromadorea). Four additional taxa (*Atractodes* sp., Cryptinae sp., Ichneumonidae sp., Pteromalidae sp.) were present in the unfiltered data but were removed by minimum sequence copy thresholds (Figure S3). The parasitoid communities associated with CSFB larvae did not significantly differ between regions of England (MGLM: Dev_4,169_ = 22.09, p = 0.599).

**Figure 5:**
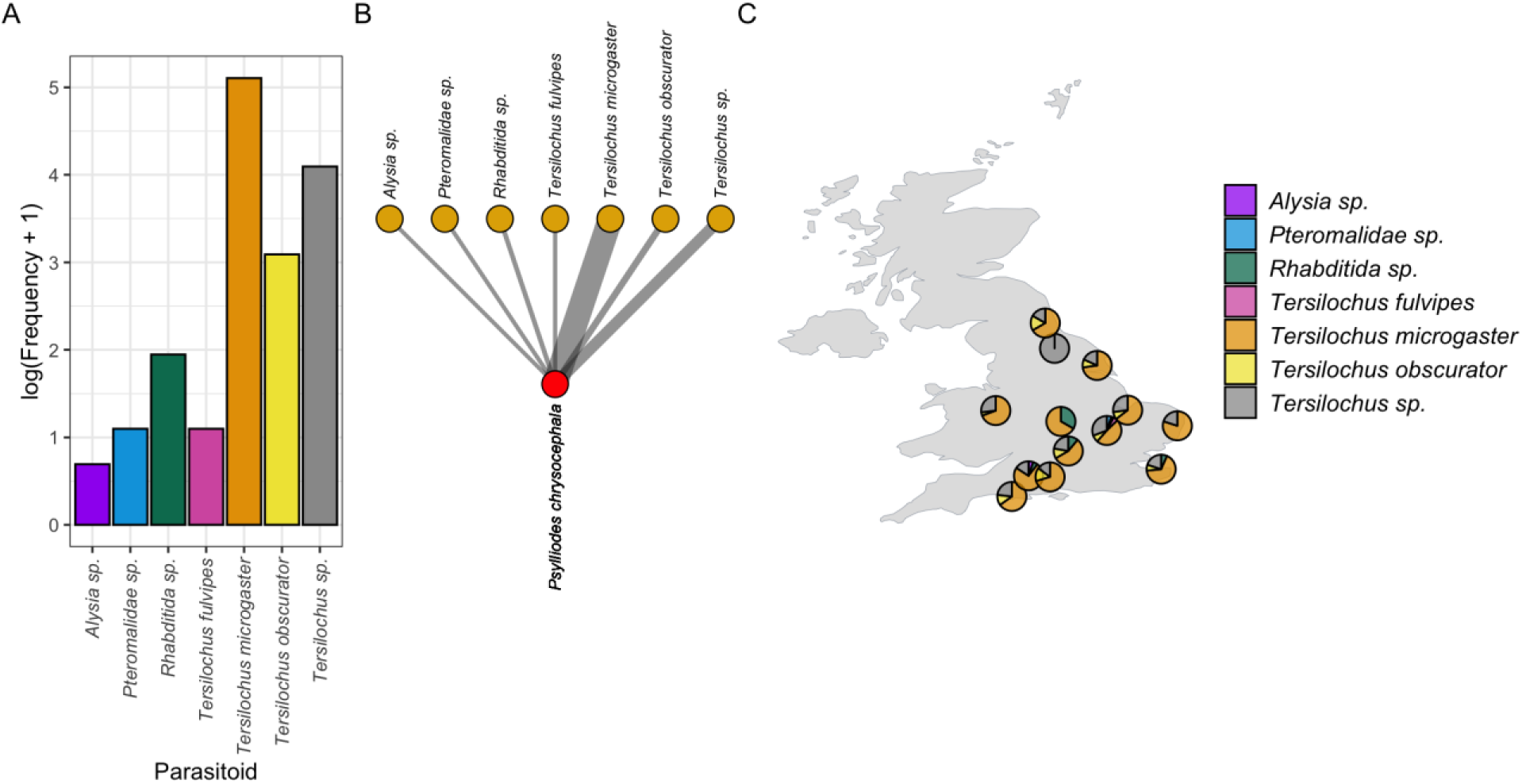
Frequency of parasitoid detections. (A), meta-network of parasitoid-cabbage stem flea beetle (CSFB) interactions across all sites in this study (B), and distribution of parasitoids detected at the sites included in this study (C). The frequency plot represents the log-scaled detection frequency for each parasitoid across the full dataset. The network represents all detected parasitoids, with link weights denoting the frequency of their detected interactions with CSFB. The filled points on the map indicate the locations from which CSFB were collected and the proportion of detected parasitoids represented by each of the detected taxa. An alternative representation for unfiltered metabarcoding data is presented in Figure S3.

## Discussion

Using a novel multi-marker metabarcoding workflow, we have identified an unexpectedly broad range of parasitoids attacking CSFB, with variation in their prevalence across regions of England. To our knowledge, this is the first study to characterise the parasitoids attacking CSFB larvae and their spatial distribution at a national scale, and the first to do so using DNA metabarcoding. Previous work has either quantified parasitism rates without identifying the species (Cerrutti et al., 2024; Serée et al., 2023) or reported individual parasitoid species within fields without assessing their distribution or parasitism rate across a wider geographic range (Barari et al., 2005; Döring et al., 2014; Ferguson et al., 2006). By integrating a novel beetle-exclusion molecular assay into a multi-marker metabarcoding study of nationwide samples, we simultaneously identified the parasitoids present, their relative frequencies, and how parasitism varies across England. This information is key for evaluating these natural enemies as conservation biocontrol agents within an IPM framework.

### Parasitoid detections and identities

The overall parasitism rate recorded in this study (11.5%) falls below that reported across Europe, where larval parasitism of CSFB varies markedly with year, region and method. In Germany, parasitism by *Tersilochus microgaster* ranges between 20% and 50% (Ulber & Wedemeyer, 2004; Ulber & Nitzsche, 2006; Döring et al., 2013), whereas UK field estimates have been substantially lower, at less than 10% (Barari et al., 2005; Ferguson et al., 2006). Recent French studies report intermediate values: Serée et al. (2023) recorded a mean of 29.55% ± 21.78 across fields (range 0–90%, with no significant difference between years), and Cerutti et al. (2024) reported parasitism of ∼12–36% (mean 24%) over 2019–2022 using high-resolution melting point analysis. The application of minimum sequence copy thresholds here removed several taxa that were detected at low read counts and reduced the parasitism rate considerably. This approach was used to limit the prevalence of false positives (Drake et al., 2022), particularly given the number of taxa detected with no prior evidence of parasitising CSFB. This approach can, however, result in false negative detections (Littleford-Colquhoun et al., 2022), potentially neglecting legitimate previously unrecorded interactions.

Our finding that *T. microgaster* was by far the most frequently detected parasitoid (64.3 % of detections), almost three times more than any other taxon, aligns with previous reports showing that *T. microgaster* is the most abundant and widespread larval parasitoid of CSFB in Europe (Barari et al., 2005; Ferguson et al., 2006b; Klingenberg & Ulber, 1994; Ulber & Nitzsche, 2006). The second most frequently detected taxon, *Tersilochus* sp., might simply represent *T. microgaster* individuals or gene copies with polymorphisms, or even sequencing errors, given the ASV-based approach used, which may have artificially reduced the dominance of *T. microgaster*. *Tersilochus obscurator* was the third most frequently detected parasitoid (8.2% of detections), with the remaining parasitoids (*T. fulvipes*, Pteromalidae sp (Hymenoptera: Chalcidoidea) and *Alysia* sp (Hymenoptera: Braconidae)) detected at a much lower frequency (< 2.4%).

To our knowledge, this is the first time that *T. obscurator* and *T. fulvipes* have been recorded parasitising CSFB, and the first record of *T. fulvipes* in the UK. *Tersilochus obscurator* is the main parasitoid of the cabbage stem weevil *Ceutorhynchus pallidactylus* (Marsham, 1802) larvae (Ulber et al., 2010). Adult *C. pallidactylus* migrate into OSR in March/June, their larvae mine the petioles and stems from May to June, completing development and leaving the plant by August (Williams, 2010). Samples for this study were collected in mid-March when *C. pallidactylus* larvae were still not present inside OSR. Similarly*, Tersilochus fulvipes* is the principal larval parasitoid of rape stem weevil *Ceutorhynchis napi* Gyllenhal, 1837 (Klingenberg & Ulber, 1994; Ulber et al., 2010). Females of *C. napi* lay their eggs into growing OSR stems in March/April, and larvae hatch within 1-2 weeks (Williams, 2010); however, neither rape stem weevil nor *T. fulvipes* have previously been recorded in the UK (Williams, 2010). Although it is not uncommon for parasitoids to attack different hosts depending on their availability, the different pest phenologies make it unlikely that the weevil hosts were present alongside the CSFB larvae.

An example of host shift or multiple host use in OSR is the case of *Stenomalina gracilis* (Walker, 1834; Pteromalidae), a parasitoid of rape stem weevil (*Ceutorhynchus napi* Gyllenhal, 1837) reported in mainland Europe that, in the United Kingdom — where the rape stem weevil is absent — instead attacks the cabbage seed weevil (*Ceutorhynchus obstrictus* (Marsham, 1802); Williams, 2010). The list of hosts a parasitoid is known to attack in the field is usually shorter than the full range of hosts it could actually develop in (Godfray, 1994). Many parasitoids lay eggs in more than one host, especially when alternative hosts can be found in the same place and may give off the same cues as the preferred host (Heimpel et al., 2003). In OSR, CSFB larvae and the stem-mining weevil larvae feed and develop inside petioles and stems, so a parasitoid searching for one could easily come across and attack the other. Also, CSFB larvae are present in the crop before weevil larvae become available (Williams, 2010). An earlier-developing host such as CSFB could therefore supply these parasitoids with hosts at the very start of the season, before its usual weevil host appears, broadening the window over which suitable hosts can be found (Godfray et al., 1994).

The two remaining detected parasitoids were Pteromalidae sp. and *Alysia* sp. The Pteromalidae are a large, worldwide family of about 418 genera and over 3,000 species, most of them in the subfamily Pteromalinae (Burks et al., 2025). Most are solitary ectoparasitoids that develop on the larvae or pupae of beetles and other insects, though some develop inside their hosts (Burks et al., 2025). Only one pteromalid, the ectoparasitoid *Trichomalus lucidus*, has been reared from CSFB before (a single female from 260 larvae, 0.3%; Nissen, 1997; Ulber & Wedemeyer, 2004). There are three pteromalid ectoparasitoids that attack cabbage seed weevil: *Trichomalus perfectus*, *Stenomalina gracilis* and *Mesopolobus morys*, all recorded in the UK (Ulber et al., 2010). *Trichomalus lucidus*, *T. perfectus* and *S. gracilis* all have COI barcodes in GenBank, so, as long as the amplicon provides sufficient resolution, species-level assignment should be possible; however, *M. morys* has no COI reference data available, so a parasitoid of this species would be resolved to family or genus. Therefore, as *M. morys* is also common in OSR and present in the UK, it is a possible identity for this detection. *Alysia*, similarly detected at a low frequency, is a large genus in the subfamily Alysiinae with 125 species worldwide, 17 of them recorded in Britain and Ireland, and all are koinobiont endoparasitoids of fly larvae (Broad et al., 2016; Sohn et al., 2023).

A further notable detection was an unidentified nematode of the order Rhabditida. This is a large and ecologically varied group containing many free-living, soil-dwelling and bacterial-feeding nematodes, as well as entomopathogenic nematodes (EPNs; De Ley, 2006). EPNs from the families *Steinernematidae* and *Heterorhabditidae* are effective biocontrol agents and have been used to control soil-dwelling larvae of leafminers, thrips, craneflies, garden chafers and various species of moth and weevils (Miles et al., 2014; Shapiro-Ilan et al., 2017). In laboratory conditions, the EPNs *Heterorhabditis bacteriophora*, *Steinernema feltiae*, *S. carpocapsae* and *S. kraussei* cause 70–85% mortality of CSFB adults (Price et al., 2023). *Steinernema feltiae,* together with a cold-active strain of *H. bacteriophora,* also reduces CSFB abundance in pot and field trials, with the third larval instar the most susceptible stage (Godina et al., 2023).

### Parasitism rates

Our results showed that, although the parasitoid community attacking CSFB did not differ between regions, the overall parasitism rate did, being significantly lower in the north. Parasitoid abundance ultimately tracks host availability over time (Godfray, 1994), and northern England has historically had both a smaller OSR area and lower CSFB pressure than southern and eastern regions (Ortega-Ramos et al., 2023). Regions with persistently low host and crop availability may therefore sustain smaller and less-established parasitoid populations than regions where the host crop is grown more widely. Differences in temperature between north and south England could also impact parasitism since the main CSFB parasitoids are active between February and April (Ulber et al., 2010), and perform best within a relatively narrow range of 14–24 °C (Johnen et al., 2010). Because northern England is cooler than the south, temperatures during this short activity window are more likely to fall below the parasitoids’ optimum, which could reduce their population sizes, shorten their period of activity, or shift their seasonal peak later than in southern regions. This gradient is, however, unlikely to be static. Larval parasitism of CSFB is known to vary considerably between years and between geographic contexts (Ferguson et al., 2006b; Ulber & Wedemeyer, 2004). Therefore, more work needs to be done to examine variation between years and to compare parasitism rates with management, landscape and weather records to disentangle the drivers of parasitism. Understanding how agri-environment measures, such as the flower-rich margins or hedgerows funded through the Sustainable Farming Incentive (SFI), may influence parasitoid populations differently along landscape and weather effects, could help tailor those measures to enhance biological control. This will require adequate investment in long-term, nationwide monitoring.

### Novel PCR primers with diagnostic potential

The novel PCR primers presented in this report exhibit a high degree of specificity toward parasitoid DNA and avoid amplification of beetle DNA. Their specificity means that they could even be used as a diagnostic tool for parasitism without downstream sequencing whilst retaining the option of sequencing where species-level resolution is desired. The theoretical taxonomic breadth of PeteBestF-BillyPrestonR also extends its applicability beyond CSFB parasitism. The same protocol could, in principle, be applied to assess parasitism in other beetle pests, such as pollen beetle (*Brassicogethes* spp.; Büchi, 2002). Importantly, this parasitoid detection is not restricted to hymenopteran parasitoids given the high prevalence of non-target groups such as Diptera in both the sequencing outputs and the *in silico* tests. This extends the potential application of these primers into biocontrol by dipteran parasitoids of pest beetles such as the Colorado potato beetle, which is parasitised by tachinid flies (Feener & Brown, 1997; López et al., 1995).

The specificity of these primers could make them suitable for adaption to loop-mediated isothermal amplification (LAMP), which has been successfully applied to rapid detection of parasitism (Ahuja & Somvanshi, 2021; Ikadai et al., 2004; Miller et al., 2026) including in field settings (Sirichaisinthop et al., 2011). If these primers were adapted for LAMP, this would form the basis of a highly accessible and user-friendly field-deployable diagnostic tool, but throughput may be limited. They could also be used in quantitative PCR (qPCR), facilitating potentially higher throughput and estimation of template DNA concentration, as demonstrated for parasitoid detection in oriental fruit flies (*Bactrocera dorsalis* (Hendel, 1912); Liang et al., 2015). Integrating these primers into a scaled workflow in which LAMP is used for rapid in-field diagnostics, followed by qPCR validation and sequencing of subsets for parasitoid identities would enable increasingly rapid, highly resolved and accurate diagnosis of parasitism. This would greatly increase the value of these primers as a tool for farmers and land managers to inform the application of insecticides according to in-field parasitism rates.

### Practical implications

Our findings have significant implications for the monitoring, study and management of parasitoid natural enemies, with particular importance for IPM practices. The PCR primers developed here can be used in a rapid diagnostic assay for parasitism. Applied directly to beetle larvae and adults, this approach could detect parasitoids much earlier than visual assessment, facilitating real-time influence on insecticide application decisions. Pesticide decision-making in OSR has traditionally ignored parasitism due to the latency that visual assessment introduces to monitoring (Barari et al., 2006). Although in the case of CSFB, detection of parasitised larvae is too late to avoid crop damage (as parasitoids do not kill the host larvae until they have completed development and enter pupation outside of the plant (Barari et al., 2005)) accounting for parasitoid prevalence in pesticide application decisions helps to safeguard parasitoid populations which will provide a greater crop protection in the long term (Farrokhzadeh et al., 2017). By extending this process to include sequencing-based identification of parasitoids, the misidentifications common in traditional approaches to parasitoid monitoring can be avoided for more accurate information on parasitoid dynamics in these pest populations (Ferguson et al., 2006a; Miller et al., 2021; Sow et al., 2019).

Understanding which parasitoids are present and which we want to promote in the field can also guide the development of bespoke landscape features, such as flower-rich margins, tailored for these species (Belz et al., 2013; Hatt et al., 2018; Jönsson et al., 2005; Skellern et al., 2023). Using the IPM framework to full effect will require knowing the host ranges of these parasitoids, and understanding their ecology and landscape needs, so that conservation strategies can be designed to support their populations. DNA metabarcoding is making this increasingly achievable by resolving host-parasitoid interactions directly (Miller et al., 2021), and network science tools already exist to turn such interaction data into practical management, for example by identifying which plants in field margins support multiple ecosystem services (Tavella et al., 2022; Windsor et al., 2021).

This approach to identifying parasitoids within beetle larvae also facilitates inclusion of parasitoids in complex interaction networks which can be used to assess more broadly the flux of ecosystem service provision in agricultural contexts (Pocock et al., 2012; Windsor et al., 2022). Parasitoids are a neglected component of such networks due to the difficulty involved in identifying them and their interactions using traditional techniques (Miller et al. 2021). High-throughput sequencing overcomes this and can even resolve complex interactions such as multiparasitism and hyperparasitism. By surveying pests and other beneficial invertebrates alongside parasitoids, networks can be constructed using a combination of direct observation and molecular data (Cuff et al., 2022). Understanding these links between pests, parasitoids and other beneficials is central to designing effective IPM, and combining molecular detection with network analysis offers a powerful framework for doing so.

## Author contributions

Patricia A. Ortega-Ramos: Conceptualization; writing – original draft; data curation; formal analysis; investigation; methodology; writing – original draft; writing – review & editing. Jordan P. Cuff: Data curation; formal analysis; validation; investigation; visualisation; methodology; writing – original draft; writing – review & editing. Darren Evans: supervision; review & editing; funding acquisition; project administration. Larissa Collins: supervision, sample administration; funding acquisition, review & editing. Alice Mauchline & Robbie Girling: supervision, funding acquisition, project administration; Samantha M. Cook: Conceptualization; supervision; funding acquisition; investigation; methodology; project administration; writing – original draft; writing – review & editing.

## Acknowledgments

This work was funded by a Rothamsted Research-University of Reading Alliance studentship to PO-R. JPC, DME and SMC were funded by the European Union’s Horizon 2020 Research and Innovation Programme as part of the EcoStack project (Grant Agreement no. 773554). PO-R and SMC are partially funded by the UK BBSRC’s Growing Health Institute Strategic Programme [BB/X010953/1]. We thank Kirsten Miller for initial discussions on this work. Provision of data is gratefully acknowledged from the Defra-funded and Fera Science Ltd managed and delivered annual Pest and Disease Survey data. For the purpose of open access, the authors have applied a Creative Commons Attribution (CC BY) licence to any Author Accepted Manuscript version arising from this submission.

## Data availability

All data are openly available via Zenodo: https://doi.org/10.5281/zenodo.23234376

## Supplementary Information

**Figure S1:**
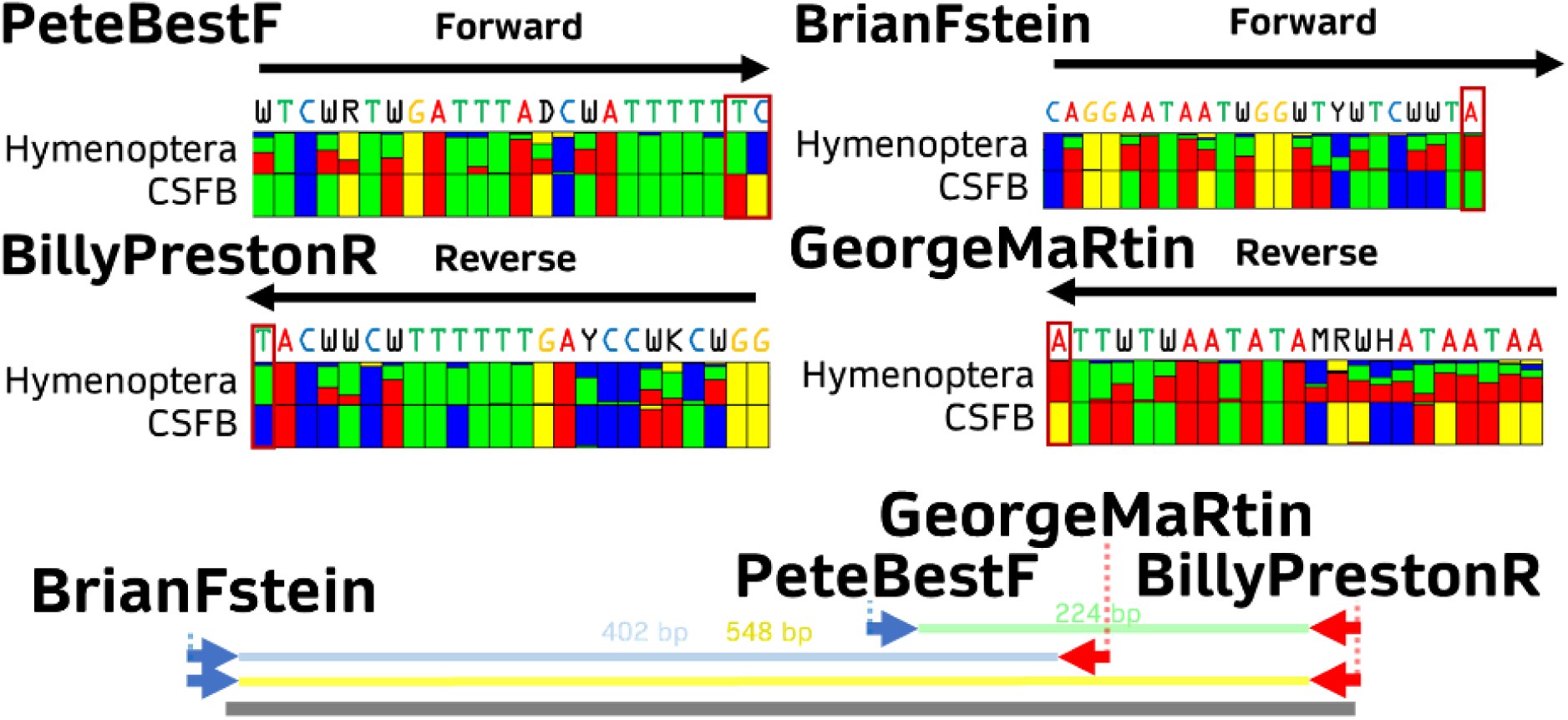
Exclusion PCR primers for the analysis of hymenopteran parasitoids in *Psylliodes chrysocephala* (cabbage stem flea beetle; CSFB). *In silico* tests indicate that the three viable pairings of these primers and one pairing with a general invertebrate primer (BerenF; Cuff et al., 2021) exhibit varying degrees of success with Hymenoptera. In all cases, however, amplification of CSFB is expected to be excluded, with variable effects on broader Coleoptera and non-target Hemiptera. The different amplicon lengths presented render this approach viable to short– or long-read sequencing.

**Figure S2:**
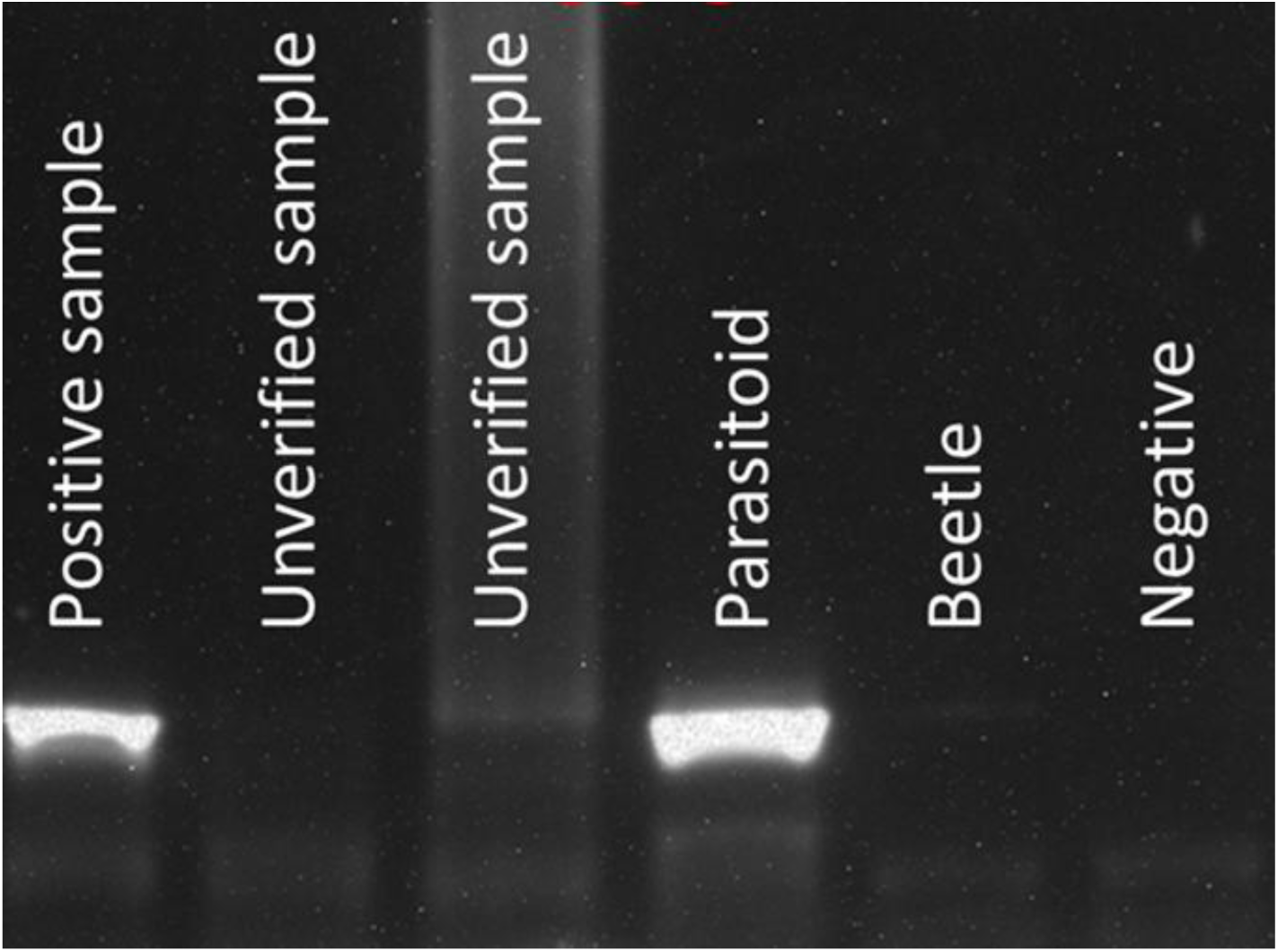
Amplification of parasitoid DNA and exclusion of beetle DNA by novel primer pair PeteBestF-BillyPrestonR. DNA was amplified from one cabbage stem flea beetle (CSFB; *Psylliodes chrysocephala*) larval sample with visually verified parasitism (‘positive sample’) and DNA extracted from an adult parasitoid (‘parasitoid’), but not from two CSFB larvae without visually obvious parasitism (‘unverified sample’) nor DNA extracted from a beetle (‘beetle’).

**Figure S3:**
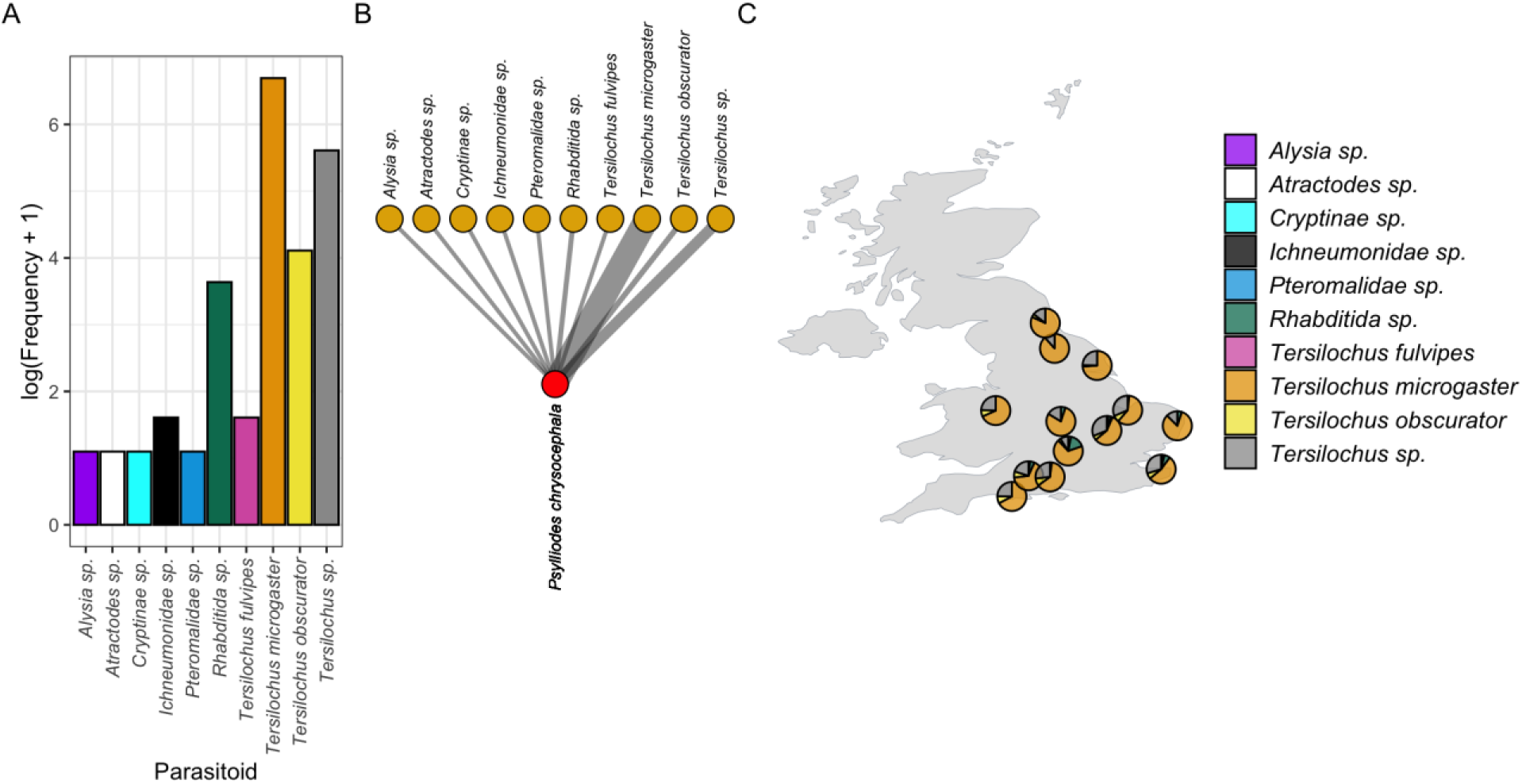
Frequency of parasitoid detections. (A), meta-network of parasitoid-cabbage stem flea beetle interactions across all sites in this study (B), and distribution of parasitoids detected at the sites included in this study (C), all for metabarcoding data to which no minimum sequence copy thresholds were applied. The frequency plot represents the log-scaled detection frequency for each parasitoid across the full dataset. The network represents all detected parasitoids, with link weights denoting the frequency of their detected interactions with CSFB. The filled points on the map indicate the locations from which CSFB were collected and the proportion of detected parasitoids represented by each of the detected taxa. An alternative representation for filtered metabarcoding data is presented in Figure 5.

## Notes

https://doi.org/10.5281/zenodo.23234376

### Competing Interest Statement

The authors have declared no competing interest.

